# A meiotic midbody structure in mouse oocytes acts as a barrier for nascent translation to ensure developmental competence

**DOI:** 10.1101/2022.11.17.516899

**Authors:** Gyu Ik Jung, Daniela Londoño-Vásquez, Sungjin Park, Ahna R. Skop, Ahmed Z. Balboula, Karen Schindler

## Abstract

Successful embryo development is dependent upon maternally deposited components. During egg formation, developmental competence is acquired through regulated translation of maternal mRNA stores. In addition, egg precursors undergo two rounds of chromosome segregation, each coupled to an asymmetric cytokinesis that produces two non-functional polar bodies. In somatic cells, cytokinesis produces two daughter cells and one midbody remnant (MBR), a signaling organelle assembled from the midbody (MB), which first appears in Telophase. MBs contain transcription and translation factors, and epigenetic modifiers. Once MBs mature to MBRs by abscission, they can be subsequently phagocytosed by another cell and influence cellular function or fate. Although the significance of MBs is elucidated in several cell types like neurons, cancer cells and stem cells, the presence and function of MBs in gametes and their roles in reproductive fitness are unknown. Here, we examined the formation and regulation of meiotic midbodies (mMB) in mouse oocytes. We find that although mouse oocyte mMBs contain analogous structures to somatic MBs, they also have a unique cap-like structure composed of the centralspindlin complex, and that cap formation depends upon an asymmetric microtubule abundance in the egg compared to the polar body. Furthermore, our results show that mMBs are translationally active ribonucleoprotein granules, supported by detection of ribosomes, polyadenylated mRNAs and nascent translation. Finally, by pharmacological and laser ablation-based approaches, we demonstrate that the mMB cap is a barrier to prevent translated products from leaving the egg and escaping into the polar body. Crucially, this barrier is critical for successful early embryonic development. Here, we document an evolutionary adaptation to the highly conserved process of cytokinesis in mouse oocytes and describe a new structure and new mechanism by which egg quality and embryonic developmental competence are regulated.

## Introduction

Oocytes, gametes derived from ovaries, undergo a maturation process that couples the completion of meiosis I with acquisition of developmental competence essential to support preimplantation embryogenesis. During meiotic maturation, the egg acquires developmental competence by rearranging organelles, degrading and translating maternal mRNAs, and erasing epigenetic modifications^1^. Importantly, after fertilization, early embryo development depends on proteins synthesized in the egg.

During meiosis I completion, oocytes segregate homologous chromosomes and undergo an asymmetric cytokinesis, releasing non-functional cells called polar bodies (PB) (Fig. 1a). In somatic cells, cytokinesis not only involves separation into daughter cells, but it also leads to the formation of a transient organelle: the midbody (MB)^2^. After cytokinesis, MBs are released extracellularly by abscission (Fig. 1a), enabling neighboring cells to phagocytose them^3-5^. These MB remnants (MBRs), when phagocytosed by cancer and stem cells, are correlated with regulation of tumorigenicity and stemness, respectively, suggesting that MBR uptake has cell type-specific effects^6^. Mammalian oocytes, which undergo asymmetric divisions, could have either an asymmetric abscission and inheritance of the MBR, or a symmetric abscission as observed in somatic cells (Fig. 1a). However, the formation of MBs and MBRs in oocytes are unknown.

**Figure 1.**
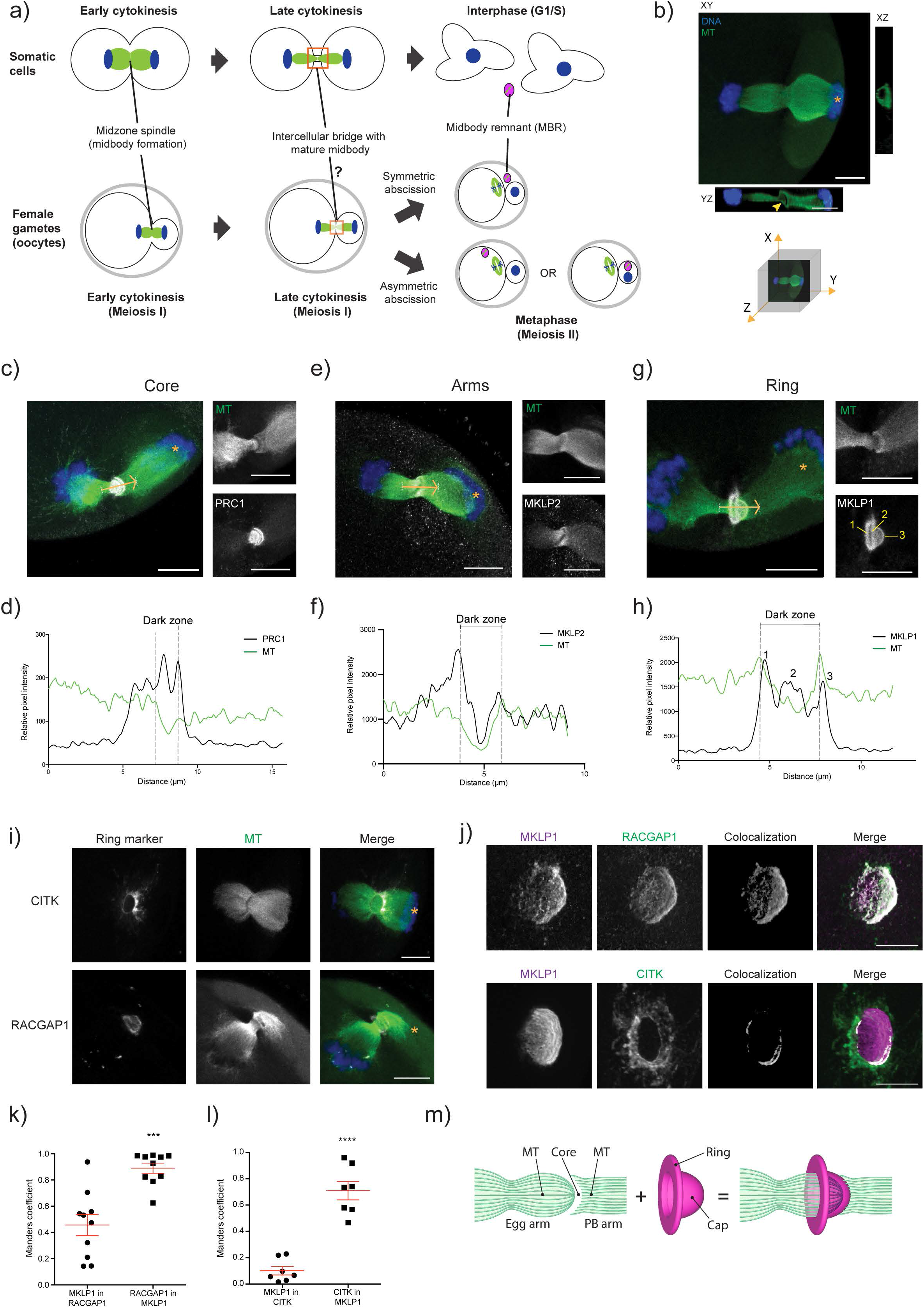
The meiotic midbody of mouse oocytes is asymmetric. a) Schematic depicting the distinct processes of mitotic cytokinesis and meiosis I cytokinesis in mammalian oocytes with the unknown outcome of midbody abscission in oocytes. b) Representative image of a mouse oocyte undergoing cytokinesis with views at XY, XZ and YZ planes. Yellow arrowhead highlights the asymmetry observed in the midzone spindle; the asterisk denotes the polar body (PB). Under confocal images is a three-dimensional coordinate system with axes depicting orientation of the different views. c, e, g) Representative z-plane projected confocal images showing localization of representative markers for the three main regions of the midbody (gray; PRC1, MKLP2 and MKLP1) relative to microtubules (green) and chromosomes (blue). Yellow arrows indicate direction of line scan plots in (d, f, h). The asterisk denotes the polar body; the numbers in the MKLP1 panel indicate the three peaks in the line scan in (h). d, f, h) Intensity line scan plots for microtubules (green) and corresponding protein (gray) of images in (c, e, g). Gray dotted lines demark the beginning and end of the midbody dark zones. i) Representative confocal images for localization of CITK (gray; top panels) and RACGAP1 (gray; bottom panels) relative to microtubules (green). The asterisk denotes the polar body. j) Representative confocal images comparing the localization of MKLP1 (magenta) with additional ring markers (gray; RACGAP1 (top panels) and CITK (bottom panels)). Signal that colocalized between the two ring components compared is shown in gray. k-l) Quantification of Manders coefficient to compare signal colocalization between (k) MKLP1 and RACGAP1, and (l) MKLP1 and CITK. Unpaired Student’s t-test, two-tailed; 10 oocytes for (k), 7 oocytes for (l). m) Schematic summarizing morphology of meiotic midbody. Scale bars = 10µm and 5µm in zoom panels; ***p < 0.001, ****p<0.0001.

The capacity of MBRs to influence cellular behavior is attributed to the proteins recruited and potentially synthesized within MBs, although little is known about how this occurs in both somatic and germ cells. Because oocytes have a limited time to produce proteins critical for successful meiosis and early embryogenesis, we hypothesized that inheritance of a translationally active meiotic MB would be critical to produce an egg developmentally competent to support early embryogenesis. Here, we report the presence of a meiotic MB (mMB) in mouse oocytes and describe a unique mMB cap-like sub-structure that arises from microtubule-led distortions, and we show that the mMB is abscised in a symmetric fashion to release a mMB remnant. We also report that the mMB is a translationally active ribonucleoprotein (RNP) granule and provide evidence that the mMB cap is a barrier for the translated products. Importantly, we demonstrate that this barrier contributes to full developmental competence of eggs by preventing maternal proteins from escaping into the first PB. Taken together, our findings highlight a mechanism by which a meiotic cell modifies mitotic machinery to provide developmental benefits for egg and embryo quality.

## Results and Discussion

Because oocytes undergo asymmetric cytokinesis (Fig. 1a), we first determined if the asymmetric division of mouse oocytes dictates morphological differences in mMBs compared to morphology of mitotic MBs. Confocal imaging of anti-tubulin-stained Telophase I-staged oocytes revealed that the microtubules at the midzone spindle are asymmetric: the spindle on the maturing oocyte (or egg) side always terminated in a ball-like structure (left) and the spindle on the PB side always terminated in a socket-like structure (right) (Fig. 1b). For ease of orientation, the egg side of all presented images will be on the left and the PB side will be on the right and labeled with an asterisk. To further investigate if the overall structure of the mMB changes because of the spindle asymmetry, we identified the three landmark regions described in mitotic MBs (ring, arms, core)^2^ by immunofluorescence to detect mitotic kinesin-like proteins 1 and 2 (MKLP1 and MKLP2) and protein regulator of cytokinesis 1 (PRC1). We examined the localization of MKLP1, MKLP2 and PRC1 proteins at meiotic stages from Metaphase I to Metaphase II, and observed dynamic localizations similar to mitotic cytokinesis^2^ (Fig. S1), which includes localization to microtubule tips at kinetochores at Metaphases I and II, and spindle midzone localization in Anaphase I. We next evaluated and compared the localization of the markers in Telophase I, the meiotic stage in which the mMB forms in mouse oocytes. PRC1 (MB core) was enriched in two disc-like structures at the spindle dark zone where the microtubule signal is absent^2^ (Fig. 1c-d), whereas MKLP2 (MB arms) colocalized with microtubules at the midzone spindle (Fig. 1e-f). These localizations are similar to observations of mitotic MBs^2^. Interestingly, centralspindlin component^7^ MKLP1 (MB ring) localization was distinct from somatic cells’ MB rings: in addition to a ring-like structure around the dark zone, which is similar to mitotic MBs, we also found a bulging, cap-like structure (cap) that surrounded the microtubules on the egg side and always protruded towards the PB (Fig. 1g-h). These localization patterns were not Telophase I-specific because we also observed the same localization patterns in Telophase II, the second asymmetric cytokinesis event in eggs (Fig. S2). We note that the cap starts to form in early Telophase I, but is distinct and fully formed in late Telophase I (data not shown). To our knowledge, this cap structure is not identified in other cell types.

To determine if the cap-and-ring structure is also observed with other MB ring markers or if this structure is unique to MKLP1, we probed Telophase I-stage oocytes for two additional markers commonly used for mitotic MB ring identification: Rac GTPase-activating protein 1 (RACGAP1), also a centralspindlin component^7^, and Citron Kinase (CIT; more commonly called CITK), the midbody kinase^8^. The images revealed that RACGAP1 localized like MKLP1 (ring + cap), whereas CITK localized only at the ring, and we did not detect a cap (Fig. 1i). We then evaluated RACGAP1 and CITK colocalization with MKLP1 by using super-resolution STED microscopy and comparing the Manders coefficients, a measure of overlap between pixels (Fig. 1j-l). The analyses indicated that there was greater colocalization of pixel signals between MKLP1 and RACGAP1 than between MKLP1 and CITK, which was expected based on their different localizations observed. These data demonstrate that mMBs have conserved structures for the arms and core as mitotic MBs, but oocytes have a modified ring that contains an additional sub-structure that bulges into the PB and consists of the centralspindlin complex that we refer here to as the cap (Fig. 1m).

We next sought to understand what drives the formation of the cap sub-structure. One of the major observable differences during cytokinesis between oocytes and most other mammalian somatic cells is that oocytes undergo an asymmetric division^9,10^ (Fig. 1a). Because of this difference, we hypothesized that the asymmetric division plays a role in mMB cap formation. To address our hypothesis, we first compressed oocytes during cytokinesis, a method which induces symmetric division and results in two daughter cells of equal size ^11^. When oocytes were forced to undergo symmetric division, we observed loss of both asymmetries (the ball/socket shape of the midzone spindle and the mMB cap) (Fig. 2a), indicating that the asymmetric cytokinetic process is involved in mMB cap formation.

**Figure 2.**
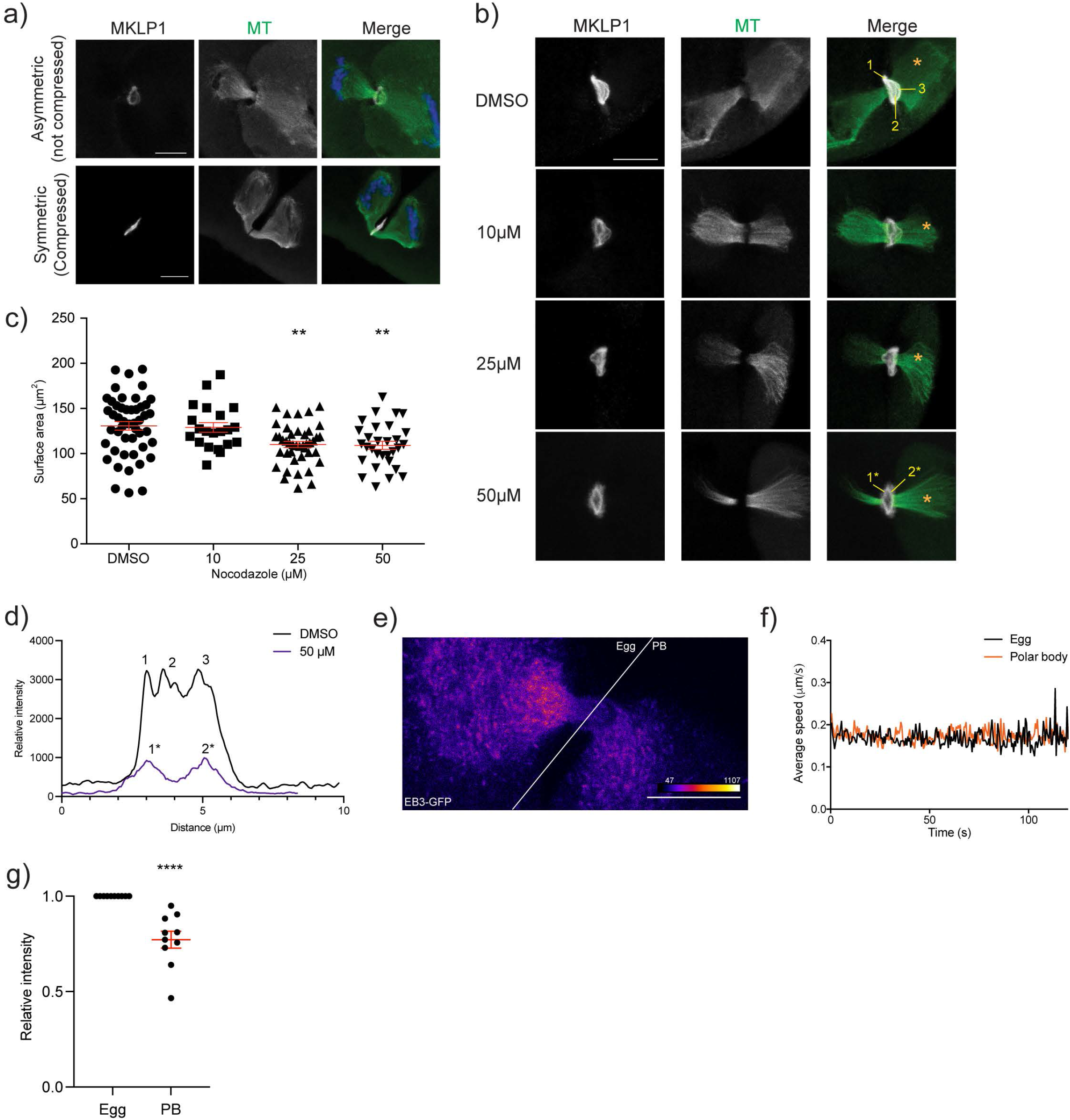
Microtubules drive distortion of meiotic MB ring and formation of meiotic midbody cap. a) Comparison of the ring structure (MKLP1, gray) and microtubules (green) when oocytes undergo asymmetric (top panels) or symmetric (bottom panels) divisions. b) Representative confocal images for nocodazole dose-dependent reduction in distortion and asymmetry of ring (gray) and midzone spindle (green). The asterisk denotes the polar body (PB) and the regions of the ring numbered are the peaks detected in the line scans in d). c) Quantification of surface area occupied by ring after nocodazole treatment. Unpaired Student’s t-test, two-tailed; 3 replicates; number of oocytes in DMSO: 48, 10 µM: 21, 25 µM: 42, 50 µM: 30; **p< 0.01. d) Plots of line scans of the intensity of MKLP1 from control and nocodazole-treated oocytes in b). The numbers reflect the numbers labeled in the corresponding images. e-g) Comparison of microtubule polymerization and dynamics during meiosis I cytokinesis by live-cell imaging of oocytes expressing EB3-GFP. e) Representative still image from live-cell confocal imaging of oocytes undergoing cytokinesis and expressing EB3-GFP. White line delineates the egg (left) and PB (right) sides. f) Average speed of EB3-GFP puncta in egg versus PB; 10 oocytes. g) Average intensity of EB3-GFP in egg versus PB. Unpaired Student’s t-test, two-tailed; 10 oocytes; ** p < 0.01, **** p< 0.0001. Scale bars = 10 µm.

Two major cellular components of cytokinesis and cell division are the midzone spindle and the actomyosin ring^12^. Because the mMB cap and ball/socket-like structure of the midzone spindle were absent when oocytes underwent symmetric division, we hypothesized that microtubules drive cap formation. To test this hypothesis, we perturbed microtubules during mMB formation using nocodazole treatment and found that increasing concentrations of this microtubule depolymerizer caused deformation of the cap at lower concentrations and complete cap regression at 50 µM, the highest concentration tested (Fig. 2b-d). Notably, at these doses, microtubules were still present, but the microtubules were symmetric because the ball-and-socket morphology disappeared. We confirmed the participation of microtubules in cap formation by live-cell imaging mMB formation in oocytes injected with *Eb3-gfp* (end-binding protein 3), a marker of a plus-end microtubules often used as an indicator of microtubule dynamics ^13^ (Fig. 2e and Video S1). By comparing microtubule polymerization speed and density between the egg and the PB sides, we found that microtubules were more abundantly polymerized on the egg side (Fig. 2e-g). These observations are in concordance with the directionality of the cap and the ball-and-socket morphology of the midzone spindle. We also tested the ability of actin to form the mMB cap. After treatment with latrunculin A, a pharmacological agent that depolymerizes actin, the cap disappeared while the ring remained (Fig. S3). But, because disruption of actin also perturbed the spindle microtubules, we cannot conclude that actin has a direct role in mMB cap formation. From these results, we concluded that microtubules distort the mMB ring and form the observed novel cap structure that is composed of centralspindlin complex proteins.

Studies on MB functions have extended beyond regulatory functions of cytokinesis, and now indicate their signaling capabilities^6^ and ribonucleoprotein (RNP) properties^14-16^. An array of proteins involved in translation, translational regulation, and RNA molecules are enriched in mitotic MBs. The enrichment of these components suggests translational capabilities within MBs and potentially explains how its uptake through phagocytosis or inheritance could regulate cellular function in a cell type-specific manner. These properties and mMB fate are unknown in oocytes. Therefore, we first investigated whether the mMB also has RNP granule characteristics, by assessing: 1) enrichment of RNA molecules, 2) increased localization of translation machinery, and 3) localized translation. By performing fluorescence *in situ* hybridization (FISH) to detect the polyadenylated (Poly-A) tail of transcripts, we found enrichment of Poly-A signal in mMBs over the background signal of RNAs in the egg cytoplasm (Fig. 3a). By immunocytochemistry, we observed that small (RPS3, RPS6, and RPS14) and large (RPL24) ribosomal subunit proteins were also enriched in mMBs (Fig. 3b). Finally, to detect nascent, active translation in the mMB, we carried out a Click chemistry-based assay that detects *L-* homopropargylglycine (HPG), a methionine-analog, that is integrated into proteins during acute incubation. Similar to mRNAs and ribosome subunit proteins, we found enrichment of nascent translation in oocyte mMBs (Fig. 3c). We confirmed the specificity of the HPG signal when we observed its decrease after treating oocytes with cycloheximide and puromycin, two translation inhibitors, and observed ∼40% reduction in HPG signal (Figs. 3c-e). We note that the HPG signal did not completely disappear. It is possible that the timing of adding the inhibitors and having translation shut down allows for some translation to occur. Alternatively, it may be difficult for chemicals to penetrate this protein dense region as also suggested by the nocodazole experiments that did not completely depolymerize microtubules.

**Figure 3.**
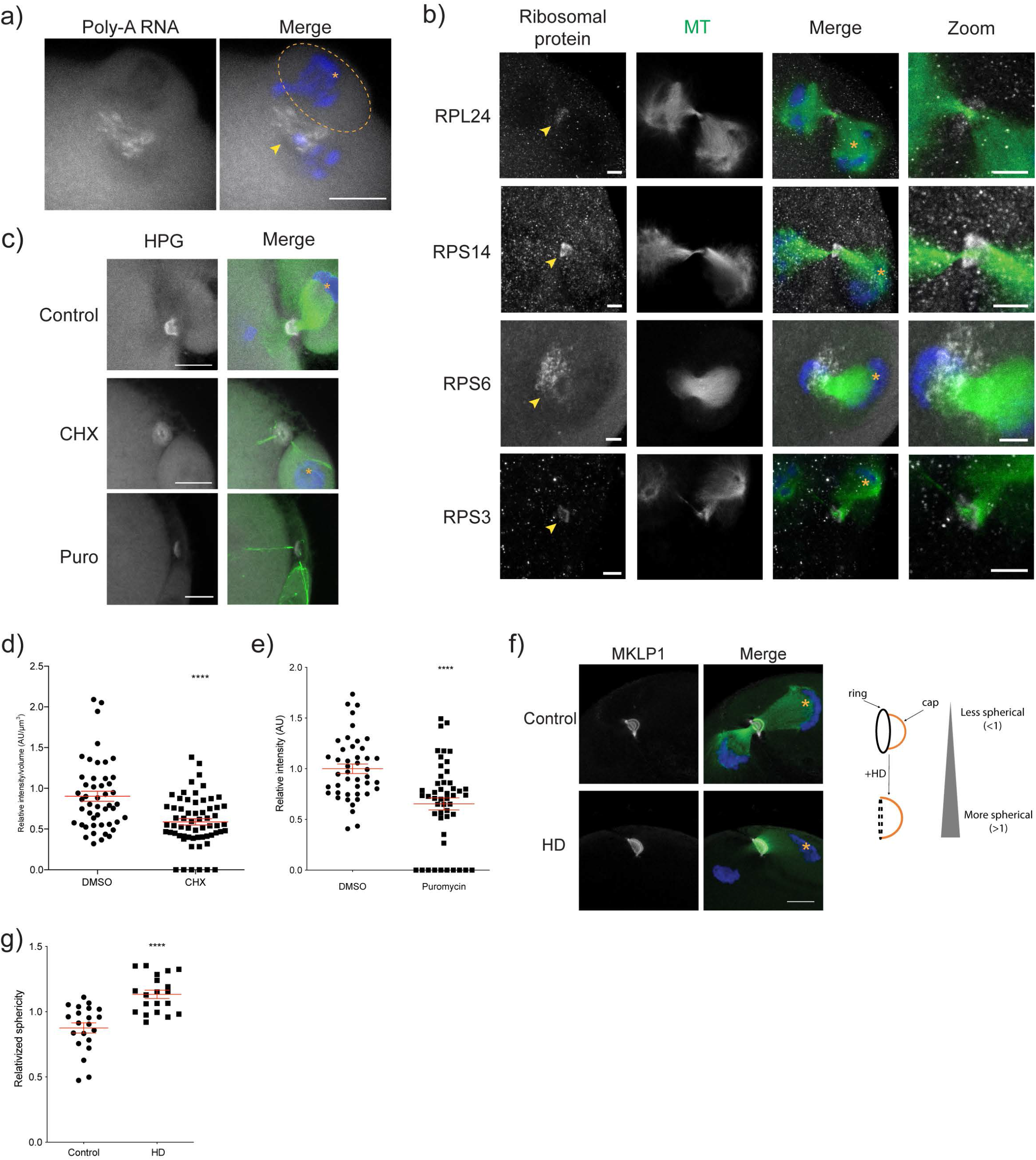
The meiotic midbody is a translationally active RNP granule. a) Confocal images showing localization of polyadenylated (Poly-A) tails of RNA molecules at the midbody region, detected by RNA FISH. The PB is encircled and denoted with an asterisk; the yellow arrowhead highlights the FISH signal enriched in the midbody region. b) Confocal images representing localization of small (RPS3, RPS6, and RPS14) and large (RPL24) ribosomal subunits (gray) at the midbody ring region relative to midzone spindle (green). The asterisk denotes the polar body (PB) and the yellow arrowhead points to the ribosomal subunit proteins. c) Representative confocal images for translational activity (gray) by Click-IT assay with and without inhibition of translation with cycloheximide (CHX) or puromycin (Puro); midzone spindle (green). HPG = homopropargylglycine; the asterisk denotes the PB. d) Quantification of translation signal at the midbody region in control versus CHX-treated cells. Unpaired Student’s t-test, two-tailed; 3 replicates; number of oocytes in DMSO: 48, CHX: 62. e) Quantification of translation signal at the midbody region in control versus puromycin-treated cells. Unpaired Student’s t-test, two-tailed; 2 replicates; number of oocytes in DMSO: 44, Puro: 49. f) Representative confocal images of midbody ring (MKLP1, gray) morphology after 1,6-hexanediol (HD) treatment. Microtubules (green) and chromosomes (blue) are also depicted. On the right, is a cartoon of the observed changes in ring and cap shapes after HD treatment. The asterisk denotes the PB. g) Quantification of sphericity as an indicator of ring morphology changes after HD treatment. Unpaired Student’s t-test, two-tailed; 3 replicates; number of oocytes in control: 21, HD: 20; ****p<0.0001. Scale bars = 10 µm and 5 µm in zoom panels.

Because RNP granules are biomolecular condensates consisting of RNA molecules and proteins that can behave like liquids^17,18^, we also tested whether mMBs are liquid-like. The aliphatic molecule 1,6-hexanediol (HD) can disrupt various phase separated, membraneless compartments^19-21^. Therefore, we evaluated the localization of MKLP1 after HD treatment of oocytes. Previous work demonstrated that 3.5% HD treatment is sufficient to disorganize a liquid-like spindle domain in mouse oocytes^22^. However, we found that this concentration did not disturb the mMB (data not shown). We therefore challenged the organization of MKLP1 with 10% HD, a concentration used in other model organisms to dissolve phase-separated structures^23^. After HD treatment, we observed that the ring, but not the cap was disrupted, suggesting that these two subregions (cap and ring) have distinct physical organizations (Fig. 3f). Consistent with phase disruption of the ring, we found that the sphericity of MKLP1, a parameter that determines how spherical a structure is, increased in the treated group because the loss of the ring structure made the overall morphology more spherical than controls (Fig. 3g). These findings support the model that the mMB is an RNP granule, consistent with our observations that there is RNA and ribosomal subunit enrichment and active translation.

To understand the mechanism by which mMBs could affect egg function, we tested two hypotheses: 1) that the mMB is abscised asymmetrically from the PB and inherited by the egg (Fig. 1a), and 2) that the cap regulates the fate of nascent translation occurring in the mMB. In somatic cells, the final stage of MB formation is abscission, or severing, of the microtubule arms, which leads to the extracellular release of the membrane-bound organelle (MBR)^24^. After release, the MBR can be internalized through phagocytosis, now called a MBsome, a necessary step for its regulatory functions^6,24^. Because we observed several asymmetries at the subcellular level in oocyte mMBs, we hypothesized that abscission of mMBs is asymmetric and is retained in the egg. To test this hypothesis, we first evaluated the recruitment of one of the endosomal sorting complexes required for transport-III (ESCRT-III) effector proteins, charged multivesicular body protein 4B (CHMP4B), at late Telophase I^25,26^. One band of CHMP4B immunoreactivity would support asymmetric abscission, whereas two parallel bands flanking the dark zone would support symmetric abscission. Here, we found CHMP4B recruited to both sides of the mMB arms and flank the dark zone (Fig. S4a), supporting a symmetric abscission model that will lead to release of a mMB remnant (mMBR). To determine if the mMB is released symmetrically, we marked mMBRs with anti-MKLP1 in Metaphase II-arrested eggs, a phase after abscission occurs. Consistent with symmetric CHMP4B localization, we found the mMBR localized in distinct foci in the perivitelline space, sandwiched between the egg and the zona pellucida and was bound by the egg and PB membranes that were marked by actin staining (Fig. S4b). From these results, we conclude that the mMB in oocytes is asymmetric in morphology but its resolution into a mMBR through abscission is symmetric as in mitosis.

We next addressed the second hypothesis, that the mMB cap controls the fate of the mMB-localized translation. One striking feature of the HPG/translation signal in mMBs was that its localization was similar to the MKLP1/RACGAP1 cap localization (Figs. 1g,i and 3c). To further evaluate the relationship between the cap and the translation signal, we imaged Telophase I-staged oocytes to detect MKLP1 and HPG Click-IT. The data indicate that the translation signal was enclosed within the cap structure, with the HPG signal enriched specifically on the egg side of the cap and absent on the PB side (Figs. 4a, S5a and Video S2). This observation led to the hypothesis that the cap is a barrier for the proteins synthesized at mMBs to remain in the egg and thereby prevent their movement into PBs. Because we previously observed that nocodazole treatment disturbs the mMB cap (Fig. 2b-d), we compared nascent translation in mMBs with an intact cap to translation when the cap was disrupted by nocodazole treatment. In contrast to control oocytes, where translation signal stopped at the MKLP1 cap signal at the egg-PB boundary, in oocytes with a disrupted cap, we saw two striking differences: 1) the translation signal no longer filled the entire space within the cap and appeared disorganized, and 2) there was HPG signal leakage into the PB (Figs. 4a and Video S3). These results suggest that the mMB cap encapsulates the translation activity and products, acting as a barrier between the egg and the PB.

**Figure 4.**
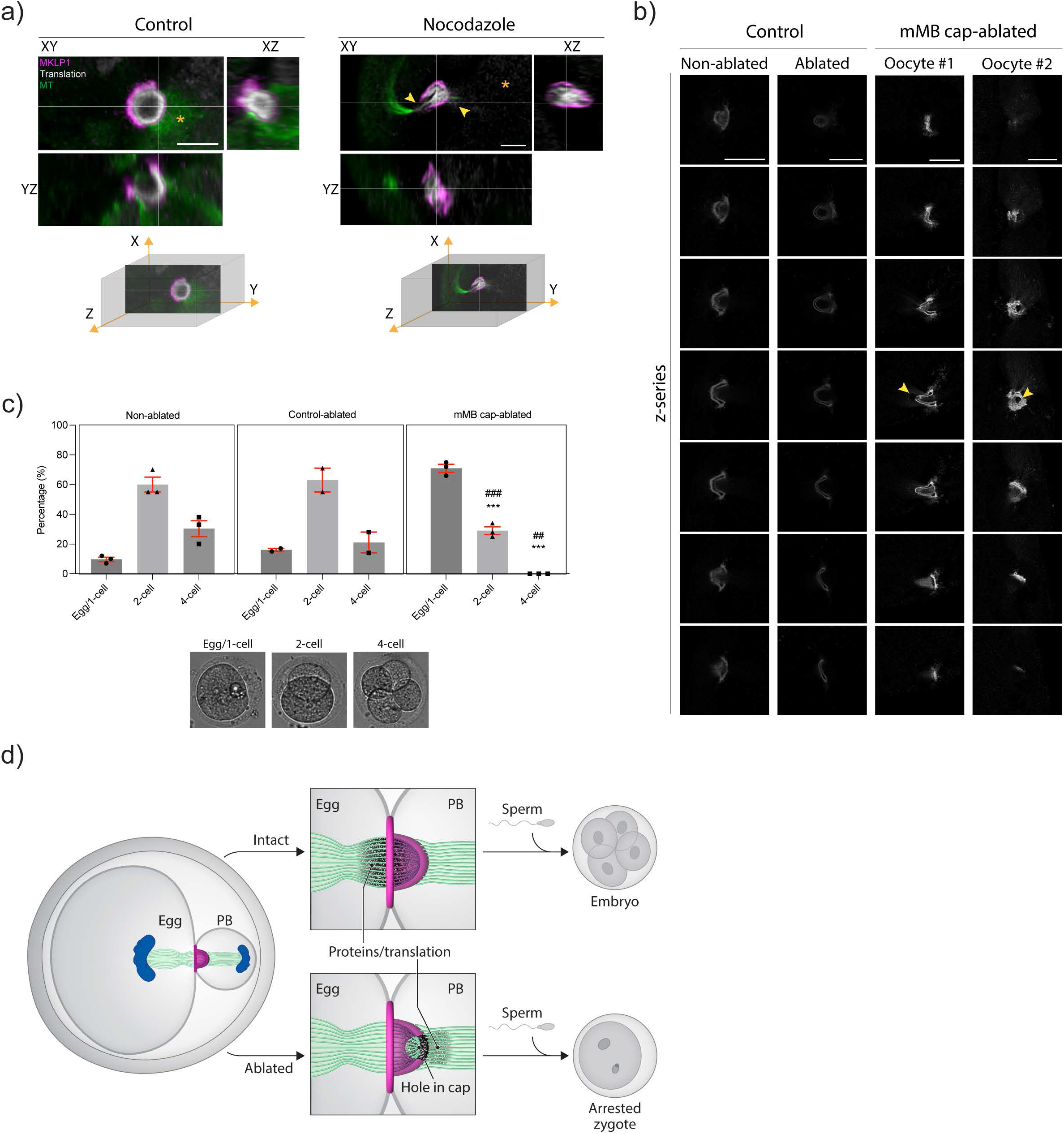
The meiotic MB cap acts as a barrier pre-abscission. a) Representative confocal images showing localization of meiotic midbody ring MKLP1 in relation to translation signal from Click-IT labeling with views at XY, XZ and YZ planes. The panels on the left are control, DMSO treated oocytes and the panels on the right are nocodazole-treated oocytes where the cap is disrupted. The asterisk denotes the polar body (PB). Under confocal images are three-dimensional coordinate systems with axes depicting orientation of the different views. b) Representative z-series confocal images of mMB caps of oocytes from non-ablated, control-ablated, and mMB cap-ablated oocytes. Two mMB cap-ablated oocytes are shown, one with a side view (oocyte #1) and one with a head-on view (oocyte #2). Arrowhead indicates where laser ablation took place. c) Images of parthenotes after ablation, activation and development *in vitro*. The graph above images quantifies the percentage of parthenotes at each developmental stage. ***p < 0.001 compared to non-ablated group; ## p < 0.01, ### p < 0.001 compared to control-ablated group. Scale bars = 10 µm. d) Model depicting structure and function of the meiotic midbody in mouse oocytes.

To test the model that the cap is a barrier and that it is important for downstream developmental competence of the egg, we used laser ablation to disrupt cap structure. Under brightfield illumination, the mMB was easily detectable because of its distinctive refraction (Fig. S5b). In Telophase I oocytes, we employed a multi-photon laser ablation (740 nm wavelength) to partially disrupt mMB cap integrity. Control ablation occurred adjacent, but not overlapping, to the mMB (Fig. S5b). We confirmed that ablation occurred in the mMB cap by detecting MKLP1 in control and cap-ablated oocytes. Control ablated oocytes had intact mMB rings and caps, whereas the cap-ablated oocytes only had MKLP1 rings and had a hole in the cap marked with anti-MKLP1 (Figs. 4b, S5b). We next parthenogenetically activated the non-ablated, control-ablated and cap-ablated Metaphase II-eggs and cultured parthenotes for two days. Approximately 80% of control parthenotes were activated and cleaved to the two- or four-cell embryonic stages. In contrast, cap-ablated parthenotes were less efficient because only ∼30% developed to the two-cell stage and no parthenotes developed to the four-cell stage (Fig. 4c). These data support the model that that mMB cap is an oocyte-specific barrier that retains the mMB translation products within the egg which later support developmental competence and preimplantation embryo development (Fig. 4d).

Our data identify mMBs in mouse oocytes, and show that they have a unique structure and RNP granule properties. The mMB cap retains nascent proteins in the egg, a function critical to subsequent embryonic development (Fig. 4c). Thus, we propose a model in which the mMB cap is an evolutionary adaptation in oocyte MBs to ensure the developmental competence of eggs after fertilization by acting as both a translation hub and as a barrier that retains maternally derived proteins in the egg (Fig. 4d).

Somatic cell MBs can regulate cellular function post-mitotically when they are phagocytosed. In *Drosophila* gonads, germ cell cysts develop from incomplete cytokinesis. The result of the incomplete cytokinesis is formation of open intracellular bridges that allow sharing of molecules and organelles, a process essential for oocyte and spermatocyte development^27-29^. A similar mechanism exists in mouse and human testes, where mitotically dividing spermatogonia undergo asymmetric cytokinesis^30,31^ and form intracellular bridges, leading to syncytia formation^32^. A key protein involved in forming these bridges is TEX14, and *Tex14* knockout male mice are infertile because spermatocytes cannot complete meiosis^33^. In the fetal mouse ovary, germ cells are also connected by intracellular bridges which later break down after birth^34^. Our data suggest that a mMB cap is a gate that closes what would otherwise become a leaky channel to keep essential proteins in the egg. In addition, our findings that mMBs are abscised symmetrically, releasing a mMBR, suggest a different mechanism to affect cell fate in embryos. We speculate that mMBR release from eggs can later act as signaling organelles during fertilization or pre-implantation embryogenesis if they are phagocytosed by the developing embryo. Further insight to the identity of the RNA transcripts and proteins in mMBs is needed to understand their roles in embryo development.

The cytoplasm of mammalian eggs sustains meiotic divisions and early embryonic development with a fixed pool of maternal transcripts that are activated and translated in a regulated fashion^35-37^. At the same time, the changes oocytes undergo throughout meiosis happen in a single cell cycle, emphasizing the need for oocytes to optimize and regulate the protein synthesis process. Spatiotemporal control of translation is found across forms of life as an energy efficient means to meet different needs during cell cycle and throughout different regions of a cell^17,38^. Oocytes lack both an interphase and an S-phase between Meiosis I and II, and they are transcriptionally silent until zygotic genome activation in embryos at the 2-cell stage in mice and 8-cell stage in humans. A recent study reports a mitochondria-dependent mode of mRNA storage in mammalian oocytes in a phase-separated, membraneless region called MARDO important for regulating translation and, consequently, proper embryo development^39^. The common theme of oocytes using membraneless organelles to control RNA storage and localization, and spindle quality^22^ points to a mechanism by which cells overcome the challenges of having a large cytoplasm. Our findings describe a third mechanism by which oocytes ensure their quality, developmental competence and potential in preparation for supporting embryogenesis. Future studies assessing mMBR fate and identification of mMBR proteins will be critical for understanding how embryos may benefit from mMBR inheritance.

## Funding

This work was supported by an NIH grant R35 GM136340 to KS. DLV and AB were supported by NIH grant R35GM142537. SP and ARS were supported by grants from NSF (MCB1158003) and NIH (GM139695-01A1).

## Author Contributions

GJ and KS, in discussion with ARS, conceived the project. GJ designed and performed experiments, data analysis, and figure preparation. DLV and AB designed and performed laser ablation experiments and analyses. GJ and KS wrote and edited the manuscript. SP, ARS, and AB provided intellectual feedback and contributed to manuscript editing.

## Competing Interest

The authors have no conflicts to disclose.

## Acknowledgements

The authors thank members of the Schindler lab for helpful discussions, Dr. Jessica Shivas for sharing her expertise and assistance with confocal and super resolution microscopy, and Dr. Di Wu for guidance with the fluorescence *in situ* hybridization protocol. The authors also thank Yi Jing Calvin Liu for his help with coding a script for analyzing live-cell imaging data and H. Adam Steinberg, owner of ArtforScience, for creating the schematic models.

## Materials and methods

### Oocyte and egg collection and culture

Sexually mature CF-1 female mice (6-10 weeks of age) were used for all experiments (Envigo, Indianapolis, IN, USA). All animals were maintained in accordance with the guidelines and policies from the Institutional Animal Use and Care Committee at Rutgers University (Protocol# 201702497) and the Animal Care Quality Assurance at the University of Missouri (Reference# 9695). Experimental procedures involving animals were approved by these regulatory bodies. Mice were housed in a room programmed for a 12-hour dark/light cycle and constant temperature, and with food and water provided *ad libitum*. Females were injected intraperitoneally with 5 I.U. of pregnant mare serum gonadotropin 48 hours prior to oocyte collection (Lee Biosolutions, Cat# 493-10). Prophase I-arrested oocytes were harvested as previously described ^40^. Briefly, cells were collected in minimal essential medium (MEM) containing 2.5 μM milrinone (Sigma-Aldrich, M4659) to prevent meiotic resumption, and cultured in Chatot, Ziomek, and Bavister (CZB) media ^41^ without milrinone in a humidified incubator programmed to 5% CO_2_ and 37° C for 11-12 hours for cytokinesis at meiosis I, or overnight for certain drug treatments.

For evaluating midbodies in meiosis II, ovulated eggs were activated with 10 mM strontium chloride (Sigma Aldrich, Cat# 25521) to induce Anaphase II onset. To collect ovulated eggs, mice were injected with human chorionic gonadotropin (hCG) (Sigma Aldrich, Cat# CG5) 48 hours after PMSG injection to stimulate ovulation of Metaphase II-arrested eggs. 14-16 hours following hCG injection, eggs were harvested from the ampulla region of the oviducts in MEM containing 3 mg/ml of hyaluronidase (Sigma Aldrich, Cat# H3506) to aid detachment of cumulus cells. Eggs were then transferred to center-well organ culture dish (Becton Dickinson, Cat# 353037) with activation media, consisting of Ca^2+^/Mg^2+^-free CZB with 10 mM of strontium chloride, and cultured in a humidified incubator programmed to 5% CO_2_ and 37°C. After 3 hours, activated eggs were cultured for 3 additional hours in KSOM + amino acids media (Sigma Aldrich, Cat# MR-106-D). For parthenogenetic activation of eggs, the activation and KSOM media were supplemented with 5 µg/ml cytochalasin D (Sigma Aldrich, Cat# C2743). Parthenogenetically activated eggs were incubated for 48 hours in KSOM + amino acids media to assess embryo cleavage rate.

For microinjection, collected oocytes were maintained arrested at prophase I with milrinone before injection to prevent nuclear disruption and after injection to allow translation of cRNAs. To induce symmetric division of oocytes, cells were compressed once they reached Metaphase I ^11^. Briefly, after culturing for 8 hours, cells were transferred to a 5-7 µl drop of CZB covered with mineral oil (Sigma Aldrich, Cat# M5310). A glass cover slip was placed on top of the media drop and pressed down on the edges to spread the media to cover the entire surface of the cover slip. The cover slip was then pressed down until oocytes flattened and the zona pellucida became indistinguishable from the cell membrane. Cells were then cultured for additional 3 hours to observe cytokinesis.

### Inhibition and disruption of mMB

To depolymerize microtubules and actin during mMB formation, oocytes were cultured in CZB for 11 hours and then transferred to media containing nocodazole (Sigma Aldrich, Cat# M1404) (0, 10, 25, and 50 µM) or latrunculin A (Cayman Chemical Company, Cat# 10010630) (0, 5, and 10 µM) in a center-well dish for 30 additional minutes.

For translation inhibition, oocytes were cultured for 9 hours prior to overnight in center-well organ dishes with CZB media supplemented with glutamine, containing either cycloheximide at 50 µg/ml (Sigma-Aldrich, Cat# C7698) or puromycin at 1 µg/ml (Sigma-Aldrich, Cat# P7255).

### Ablation of mMB cap by laser ablation

Prophase I-arrested oocytes were cultured *in vitro* in milrinone-free CZB medium supplemented with 100 nM SiR-tubulin (Cytoskeleton #NC0958386) in a humidified, microenvironmental chamber (5% CO_2_ and 37° C) equipped to a Leica TCP SP8 inverted microscope. After culturing cells for 11 hours, mMB caps were partially ablated using a multi-photon laser as previously described ^42^. In brief, a 4µm^2^ square region of interest within the mMB cap was exposed to a 740 nm wavelength and 60-70 mW power laser beam at the sample plane. For control-ablated oocytes, the cytoplasmic region adjacent to the mMB was exposed to the same protocol. A subset of cap-ablated, control-ablated and non-ablated oocytes were fixed and immunostained with MKLP1 antibody to assess the efficiency of laser ablation and mMB cap disruption.

### Immunofluorescence

Following meiotic maturation, oocytes or activated eggs were fixed in various conditions to detect localization of proteins. For detection of PRC1 (Proteintech, 15617-1-AP, 1:100), CIT-K (BD Biosciences, 611376, 1:100), RACGAP1 (Santa Cruz, sc-271110, 1:50), MKLP1 (Novus Biologicals, NBP2-56923, 1:100), and MKLP2 (Proteintech, 67190-1, 1:100), oocytes were fixed in 2% PFA in phosphate-buffered saline (PBS) for 20 minutes at room temperature. For detection of RPS3 (Cell Signaling Technology, 2579S, 1:30), RPS6 (Santa Cruz, sc-74459, 1:30), RPS14 (Proteintech, 16683-1-AP, 1:30), and RPL24 (ThermoFisher, PA5-62450, 1:30), oocytes were fixed in cold methanol (Sigma Aldrich, Cat# A452-4) for 10 minutes. For detection of CHMP4B (Proteintech, 13683-1-AP, 1:30), zona pellucida were removed from oocytes by brief treatment with acidic Tyrode’s solution (Sigma Aldrich, Cat# MR-004-D) and fixed with 2% PFA in PBS for 20 minutes at room temperature. After fixation, oocytes were incubated in blocking buffer (0.3% BSA and 0.01% Tween in PBS) for at least 10 minutes before proceeding. For permeabilization, oocytes were incubated in PBS containing 0.2% Triton-X for 20 minutes and blocked with blocking buffer for 10 minutes. For RPS3, RPS6, RPS14, RPL24, RACGAP1, and CHMP4B, cells were incubated overnight at 4° C with primary antibody. For all other proteins, primary incubation was performed for 1 hour at room temperature. For secondary antibody incubation, oocytes were incubated for 1 hour in a dark, humidified chamber at room temperature. Both antibody incubations were followed by three washes in blocking solution, 10 minutes each. After the last wash, oocytes were mounted in 10 µl of Vectashield (Vector Laboratories, Cat# H-1000) containing 4,6-Diamidino-2-Phenylindole, Dihydrochloride (DAPI) (Life Technologies, Cat# D1306; 1:170) for confocal microscopy.

For super-resolution microscopy using the tau-3X STED module from Leica Biosystems, the same steps as the ones described above for confocal microscopy were followed except for the following changes: 1) antibody concentrations were doubled for primary antibodies and 2) after the third wash following secondary antibody incubation, cells were mounted in 10 µl of EMS glycerol mounting medium with DABCO (EMS, Cat# 17989-10).

### RNA *in situ* hybridization

To detect RNA molecules, fluorescence *in situ* hybridization (FISH) against the poly-A tail of transcripts was performed using an oligo-dT probe that consists of 21 thymine nucleotides with a 3’ modification of a fluorophore as described ^43^. Briefly, oocytes were fixed in increasing volumes of methanol-free 4% formaldehyde diluted in RNase-free 1X PBS at 37° C for 45 minutes. Oocytes were then dehydrated in increasing concentrations of methanol and stored at −20° C until further processing. Oocytes were prepared for hybridization by washing through 1X PBS with 0.1% Tween-20 (PBT), followed by 10% formamide and 2X SSC in nuclease-free water (Wash A). For the hybridization reaction, oocytes were incubated in a 10% formamide, 2X SSC and 10% dextran sulfate solution in nuclease-free water with 12.5 µM of the probe overnight at 37°C. After hybridization, samples were rinsed through several volumes of fresh, pre-warmed Wash A and PBT before mounting on 10 µl of Vectashield with DAPI for imaging.

### Nascent protein detection assay

Translation activity at the midbody was assessed by detecting protein synthesis level using an *L*-HPG-translation kit (ThermoFisher, Cat# C10429) as previously described ^44^. In summary, oocytes were collected and matured for 11.5 hours, then transferred to DMEM medium lacking methionine (ThermoFisher, Cat# 42-360-032) and containing HPG diluted 1:50 for 30-45 minutes, followed by fixation with 2% PFA in PBS for 20 minutes at room temperature and subsequent detection of HPG signal by immunofluorescence.

### Image acquisition and live-cell imaging

Confocal and super-resolution images were acquired using a Leica SP8 confocal microscope with Lightning module equipped with a 40X, 1.30NA oil-immersion objective. Super-resolution STED images were acquired using a Leica SP8 confocal microscope with Tau-STED module equipped with a 93X, 1.30NA glycerol-immersion objective. For each image, optical z-sections were obtained using 0.5-1 µm step-size with a zoom factor of 2.5-6. Oocytes from experiments involving comparison of intensities or stages were processed on the same day and imaged maintaining laser settings equal across samples.

Live-cell confocal image acquisition was performed using a Leica SP8 confocal microscope system with a 40X, 1.30NA oil-immersion objective, equipped with a heated, humidified stage top incubator with 5% CO_2_ and 37° C (Tokai Hit, STX stage top incubator). To observe progression through cytokinesis, images of oocytes were acquired every 20 minutes with 15 optical sections across the entire thickness of each oocyte at 1024×1024-pixel image resolution and 600 Hz acquisition speed. For EB3-GFP tracking during cytokinesis, images were taken every 0.5 second at a single plane at 1024×512-pixel image resolution and 1000 Hz acquisition speed.

### Cloning and cRNA preparation

To generate cRNA of *Eb3-egfp* ^*45*^, the plasmid was linearized and transcribed *in vitro* using mMessage mMachine T7 kit (Ambion, Cat# AM1344) according to manufacturer’s protocol.

cRNA was purified using SeraMag Speedbead (Sigma Aldrich, Cat# GE65152105050250) nucleotide purification method previously described ^46^. Briefly, *in vitro* transcription reaction solution was brought up to 150 µl and mixed with 100 µl of magnetic beads and let stand for 5 minutes. Beads were then pelleted using a magnetic stand and washed with 80% ethanol. cRNA was eluted using 20 µl nuclease-free water and stored at −80° C.

### Image analysis and quantification

All images and videos were analyzed and quantified using Imaris software (Bitplane, Oxford Instrument Company) and Fiji ^47^. Quantification of sphericity, volume, and intensity were performed by creating a region of interest (ROI) with the “surfaces” tool in Imaris. To determine ROI, threshold of signal was determined from control groups and applied in treatment groups. For co-localization analyses, the “co-localization analysis” tool in Imaris was used. For EB3-GFP speed tracking, videos were processed by Gaussian filter blend and background subtraction. Individual puncta were then determined using the “spots” tool and filtering for spots that could be tracked in at least 3 continuous frames. For EB3-GFP intensity measurements, the first frame of each video was used to compare the intensity of the egg side to the PB side. The dark zone was used as a reference to distinguish the egg and the PB and mark ROIs.

### Statistical analysis

As indicated in the figure legends, one-way ANOVA and unpaired Student’s t-test analyses were performed to examine statistical differences between groups using GraphPad Prism software. p<0.05 was considered statistically significant. All error bars shown reflect standard errors of means.

## Figure Legends

**Figure S1. Midbody protein localization during Meiosis I and at Metaphase of Meiosis II**. a-c) Representative confocal images showing the localization of MKLP1 (a), PRC1 (b), and MLKP2 (c) at Metaphase I, Anaphase I, Telophase I, and Metaphase II; microtubules (gray/green in merge) and DAPI (blue). Scale bars = 10µm.

**Figure S2. Localization of midbody markers in Telophase of meiosis II**. Representative confocal images showing the localization of MKLP1, MKLP2, PRC1, and CITK (gray) relative to midzone spindle (green) in midbody from activated eggs; DAPI in blue. Scale bars = 10 µm.

**Figure S3. Latrunculin A treatment for actin depolymerization**. a-b) Representative confocal images showing the morphology of MKLP1 (gray) relative to midzone spindle (a) and actin (b) (green) in midbodies from oocytes treated with DMSO, or 5 µM and 10 µM of latrunculin A. c) Quantification of percentages of normal and abnormal midbodies from oocytes treated with DMSO, or 5 µM and 10 µM latrunculin A.

**Figure S4. Abscission of the meiotic midbody is symmetric**. a) Representative confocal images showing localization of CHMP4B (gray) relative to microtubules (green) and to the MB region in early Telophase I (top panels), Late Telophase I (middle panels), and Metaphase II (bottom panels). Arrowheads on CHMP4B and Zoom panels indicate regions of CHMP4B enrichment. b) Representative confocal images of MKLP1 (gray) and cell boundaries (actin, green) in Metaphase II. On the left, whole egg image with membrane delineated with dotted, white circle is shown. Orange square indicates the region shown in the zoom in images on the right side with XY, YZ, and XZ views with a three-dimensional coordinate system with axes depicting orientation of the different views. Scale bars = 5 µm, 10 µm, and 15 µm.

**Figure S5. Detection of meiotic midbody and confirmation of laser ablation**. a) Representative 3D reconstruction from confocal images showing translation localization (gray) labeled with HPG relative to MKLP1 cap (magenta). b) Representative confocal images showing microtubules labeled with SiR-Tubulin and brightfield images of Telophase I-oocytes. The area of ablation is marked with green foci and the zone after ablation is marked with a white square. The dotted white lines outline the oocyte and the polar body. Scale bars = 2 µm and 10 µm.

## Notes

### Competing Interest Statement

The authors have declared no competing interest.

## References

1 Sha, Q. Q., Zhang, J. & Fan, H. Y. A story of birth and death: mRNA translation and clearance at the onset of maternal-to-zygotic transition in mammalsdagger. Biol Reprod 101, 579–590 (2019). https://doi.org:10.1093/biolre/ioz012

2 Hu, C. K., Coughlin, M. & Mitchison, T. J. Midbody assembly and its regulation during cytokinesis. Mol Biol Cell 23, 1024–1034 (2012). https://doi.org:10.1091/mbc.E11-08-0721

3 Crowell, E. F., Gaffuri, A. L., Gayraud-Morel, B., Tajbakhsh, S. & Echard, A. Engulfment of the midbody remnant after cytokinesis in mammalian cells. J Cell Sci 127, 3840–3851 (2014). https://doi.org:10.1242/jcs.154732

4 Kuo, T. C. et al. Midbody accumulation through evasion of autophagy contributes to cellular reprogramming and tumorigenicity. Nat Cell Biol 13, 1214–1223 (2011). https://doi.org:10.1038/ncb2332

5 Peterman, E. et al. The post-abscission midbody is an intracellular signaling organelle that regulates cell proliferation. Nat Commun 10, 3181 (2019). https://doi.org:10.1038/s41467-019-10871-0

6 Farmer, T. & Prekeris, R. New signaling kid on the block: the role of the postmitotic midbody in polarity, stemness, and proliferation. Mol Biol Cell 33 (2022). https://doi.org:10.1091/mbc.E21-06-0288

7 White, E. A. & Glotzer, M. Centralspindlin: at the heart of cytokinesis. Cytoskeleton (Hoboken) 69, 882–892 (2012). https://doi.org:10.1002/cm.21065

8 Bassi, Z. I., Audusseau, M., Riparbelli, M. G., Callaini, G. & D’Avino, P. P. Citron kinase controls a molecular network required for midbody formation in cytokinesis. Proc Natl Acad Sci U S A 110, 9782–9787 (2013). https://doi.org:10.1073/pnas.1301328110

9 Green, R. A., Paluch, E. & Oegema, K. Cytokinesis in animal cells. Annu Rev Cell Dev Biol 28, 29–58 (2012). https://doi.org:10.1146/annurev-cellbio-101011-155718

10 Chaigne, A., Terret, M. E. & Verlhac, M. H. Asymmetries and Symmetries in the Mouse Oocyte and Zygote. Results Probl Cell Differ 61, 285–299 (2017). https://doi.org:10.1007/978-3-319-53150-2_13

11 Otsuki, J. et al. Symmetrical division of mouse oocytes during meiotic maturation can lead to the development of twin embryos that amalgamate to form a chimeric hermaphrodite. Hum Reprod 27, 380–387 (2012). https://doi.org:10.1093/humrep/der408

12 Normand, G. & King, R. W. Understanding cytokinesis failure. Adv Exp Med Biol 676, 27–55 (2010). https://doi.org:10.1007/978-1-4419-6199-0_3

13 Stepanova, T. et al. Visualization of microtubule growth in cultured neurons via the use of EB3-GFP (end-binding protein 3-green fluorescent protein). J Neurosci 23, 2655–2664 (2003).

14 Bonner, M. K. et al. Mitotic spindle proteomics in Chinese hamster ovary cells. PLoS One 6, e20489 (2011). https://doi.org:10.1371/journal.pone.0020489

15 Capalbo, L. et al. The midbody interactome reveals unexpected roles for PP1 phosphatases in cytokinesis. Nat Commun 10, 4513 (2019). https://doi.org:10.1038/s41467-019-12507-9

16 Skop, A. R., Liu, H., Yates, J., 3rd, Meyer, B. J. & Heald, R. Dissection of the mammalian midbody proteome reveals conserved cytokinesis mechanisms. Science 305, 61–66 (2004). https://doi.org:10.1126/science.1097931

17 Das, S., Vera, M., Gandin, V., Singer, R. H. & Tutucci, E. Intracellular mRNA transport and localized translation. Nat Rev Mol Cell Biol 22, 483–504 (2021). https://doi.org:10.1038/s41580-021-00356-8

18 Brangwynne, C. P. et al. Germline P granules are liquid droplets that localize by controlled dissolution/condensation. Science 324, 1729–1732 (2009). https://doi.org:10.1126/science.1172046

19 Yang, C., Dominique, G. M., Champion, M. M. & Huber, P. W. Remnants of the Balbiani body are required for formation of RNA transport granules in Xenopus oocytes. iScience 25, 103878 (2022). https://doi.org:10.1016/j.isci.2022.103878

20 Geiger, F. et al. Liquid-liquid phase separation underpins the formation of replication factories in rotaviruses. EMBO J 40, e107711 (2021). https://doi.org:10.15252/embj.2021107711

21 Agote-Aran, A. et al. Spatial control of nucleoporin condensation by fragile X-related proteins. EMBO J 39, e104467 (2020). https://doi.org:10.15252/embj.2020104467

22 So, C. et al. A liquid-like spindle domain promotes acentrosomal spindle assembly in mammalian oocytes. Science 364 (2019). https://doi.org:10.1126/science.aat9557

23 Ding, D. Q. et al. Chromosome-associated RNA-protein complexes promote pairing of homologous chromosomes during meiosis in Schizosaccharomyces pombe. Nat Commun 10, 5598 (2019). https://doi.org:10.1038/s41467-019-13609-0

24 Peterman, E. & Prekeris, R. The postmitotic midbody: Regulating polarity, stemness, and proliferation. J Cell Biol 218, 3903–3911 (2019). https://doi.org:10.1083/jcb.201906148

25 Petsalaki, E. & Zachos, G. The Abscission Checkpoint: A Guardian of Chromosomal Stability. Cells 10 (2021). https://doi.org:10.3390/cells10123350

26 Renshaw, M. J., Liu, J., Lavoie, B. D. & Wilde, A. Anillin-dependent organization of septin filaments promotes intercellular bridge elongation and Chmp4B targeting to the abscission site. Open Biol 4, 130190 (2014). https://doi.org:10.1098/rsob.130190

27 Greenbaum, M. P., Iwamori, T., Buchold, G. M. & Matzuk, M. M. Germ cell intercellular bridges. Cold Spring Harb Perspect Biol 3, a005850 (2011). https://doi.org:10.1101/cshperspect.a005850

28 Huynh, J. R. & St Johnston, D. The origin of asymmetry: early polarisation of the Drosophila germline cyst and oocyte. Curr Biol 14, R438–449 (2004). https://doi.org:10.1016/j.cub.2004.05.040

29 Lu, K., Jensen, L., Lei, L. & Yamashita, Y. M. Stay Connected: A Germ Cell Strategy. Trends Genet 33, 971–978 (2017). https://doi.org:10.1016/j.tig.2017.09.001

30 Huckins, C. The spermatogonial stem cell population in adult rats. I. Their morphology, proliferation and maturation. Anat Rec 169, 533–557 (1971). https://doi.org:10.1002/ar.1091690306

31 de Rooij, D. G. The nature and dynamics of spermatogonial stem cells. Development 144, 3022–3030 (2017). https://doi.org:10.1242/dev.146571

32 Fawcett, D. W., Ito, S. & Slautterback, D. The occurrence of intercellular bridges in groups of cells exhibiting synchronous differentiation. J Biophys Biochem Cytol 5, 453–460 (1959). https://doi.org:10.1083/jcb.5.3.453

33 Greenbaum, M. P. et al. TEX14 is essential for intercellular bridges and fertility in male mice. Proc Natl Acad Sci U S A 103, 4982–4987 (2006). https://doi.org:10.1073/pnas.0505123103

34 Pepling, M. E. & Spradling, A. C. Mouse ovarian germ cell cysts undergo programmed breakdown to form primordial follicles. Dev Biol 234, 339–351 (2001). https://doi.org:10.1006/dbio.2001.0269

35 Luong, X. G., Daldello, E. M., Rajkovic, G., Yang, C. R. & Conti, M. Genome-wide analysis reveals a switch in the translational program upon oocyte meiotic resumption. Nucleic Acids Res 48, 3257–3276 (2020). https://doi.org:10.1093/nar/gkaa010

36 Yang, C. R. et al. The RNA-binding protein DAZL functions as repressor and activator of mRNA translation during oocyte maturation. Nat Commun 11, 1399 (2020). https://doi.org:10.1038/s41467-020-15209-9

37 Chen, J. et al. Genome-wide analysis of translation reveals a critical role for deleted in azoospermia-like (Dazl) at the oocyte-to-zygote transition. Genes Dev 25, 755–766 (2011). https://doi.org:10.1101/gad.2028911

38 Chaigne, A. & Brunet, T. Incomplete abscission and cytoplasmic bridges in the evolution of eukaryotic multicellularity. Curr Biol 32, R385–R397 (2022). https://doi.org:10.1016/j.cub.2022.03.021

39 Cheng, S. et al. Mammalian oocytes store mRNAs in a mitochondria-associated membraneless compartment. Science 378, eabq4835 (2022). https://doi.org:10.1126/science.abq4835

40 Blengini, C. S. & Schindler, K. Immunofluorescence Technique to Detect Subcellular Structures Critical to Oocyte Maturation. Methods Mol Biol 1818, 67–76 (2018). https://doi.org:10.1007/978-1-4939-8603-3_8

41 Chatot, C. L., Ziomek, C. A., Bavister, B. D., Lewis, J. L. & Torres, I. An improved culture medium supports development of random-bred 1-cell mouse embryos in vitro. J Reprod Fertil 86, 679–688 (1989). https://doi.org:10.1530/jrf.0.0860679

42 Londono-Vasquez, D., Rodriguez-Lukey, K., Behura, S. K. & Balboula, A. Z. Microtubule organizing centers regulate spindle positioning in mouse oocytes. Dev Cell 57, 197–211 e193 (2022). https://doi.org:10.1016/j.devcel.2021.12.011

43 Wu, D. & Dean, J. EXOSC10 sculpts the transcriptome during the growth-to-maturation transition in mouse oocytes. Nucleic Acids Res 48, 5349–5365 (2020). https://doi.org:10.1093/nar/gkaa249

44 Aboelenain, M. & Schindler, K. Aurora kinase B inhibits aurora kinase A to control maternal mRNA translation in mouse oocytes. Development 148 (2021). https://doi.org:10.1242/dev.199560

45 Schuh, M. & Ellenberg, J. Self-organization of MTOCs replaces centrosome function during acentrosomal spindle assembly in live mouse oocytes. Cell 130, 484–498 (2007). https://doi.org:10.1016/j.cell.2007.06.025

46 Modzelewski, A. J. et al. Efficient mouse genome engineering by CRISPR-EZ technology. Nat Protoc 13, 1253–1274 (2018). https://doi.org:10.1038/nprot.2018.012

47 Schindelin, J. et al. Fiji: an open-source platform for biological-image analysis. Nat Methods 9, 676–682 (2012). https://doi.org:10.1038/nmeth.2019

