## Supplementary figures and images for "A meiotic midbody structure in mouse oocytes acts as a barrier for nascent translation to ensure developmental competence"

### Fig. S1

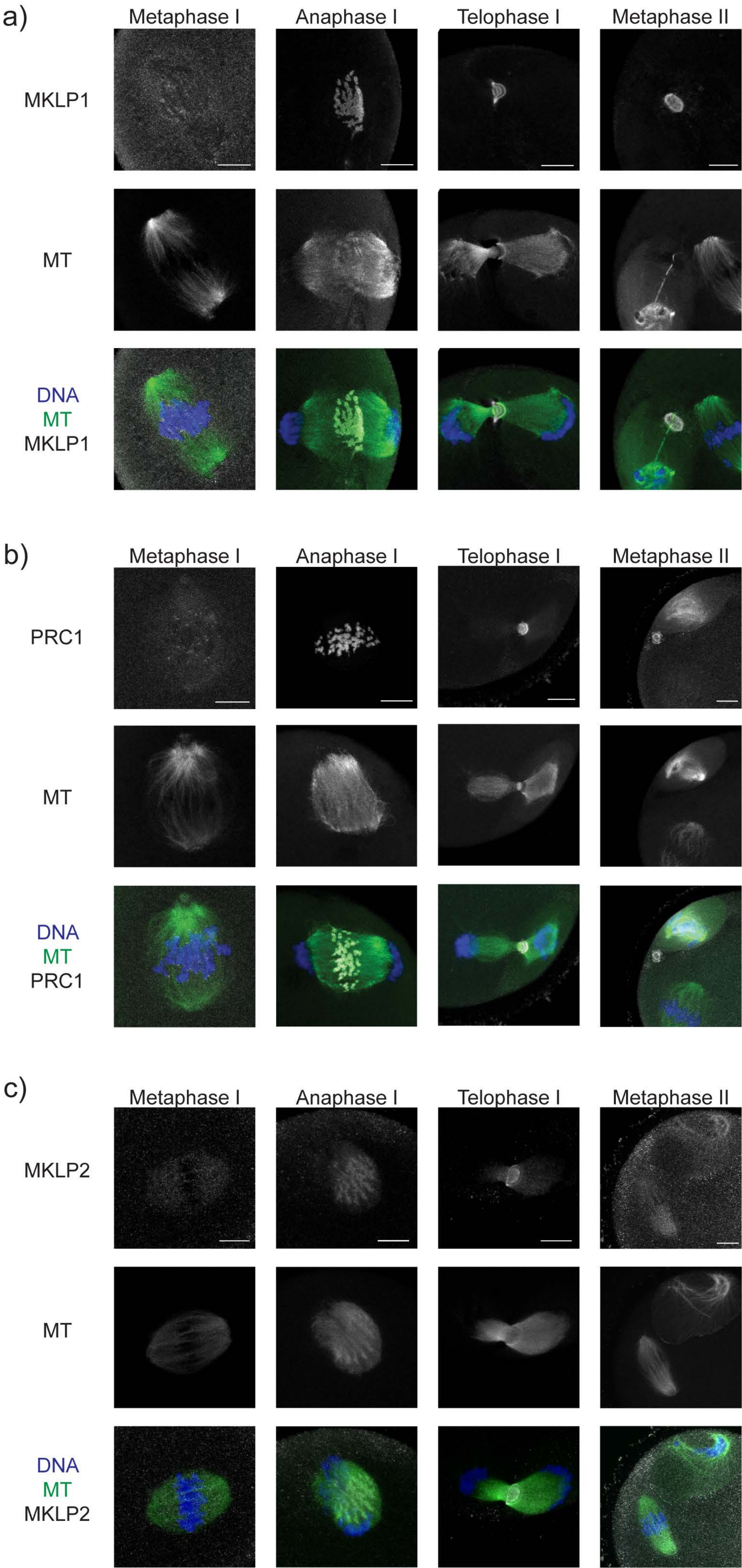

### Fig. S2

#### Figure S2

a)

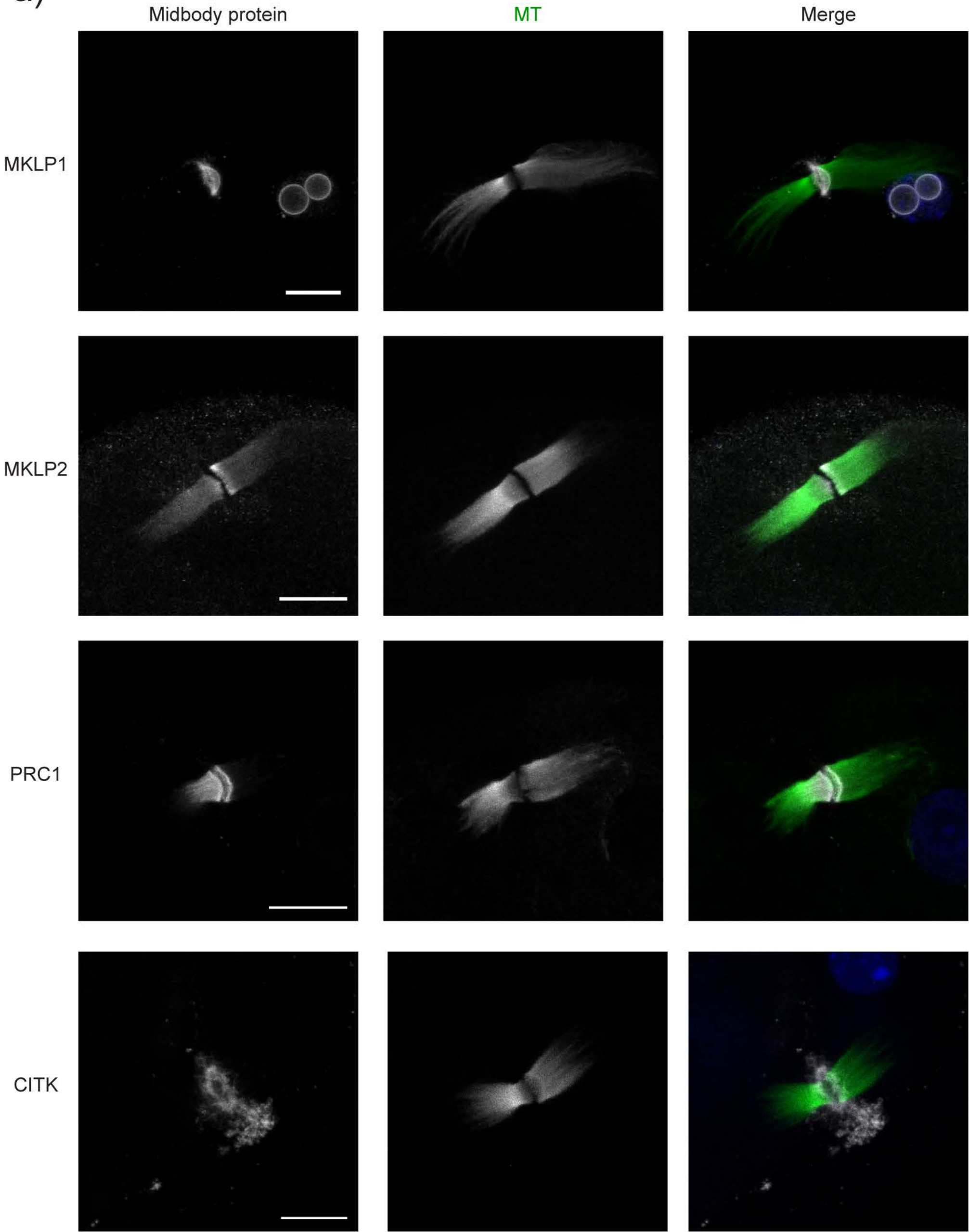

### Fig. S3

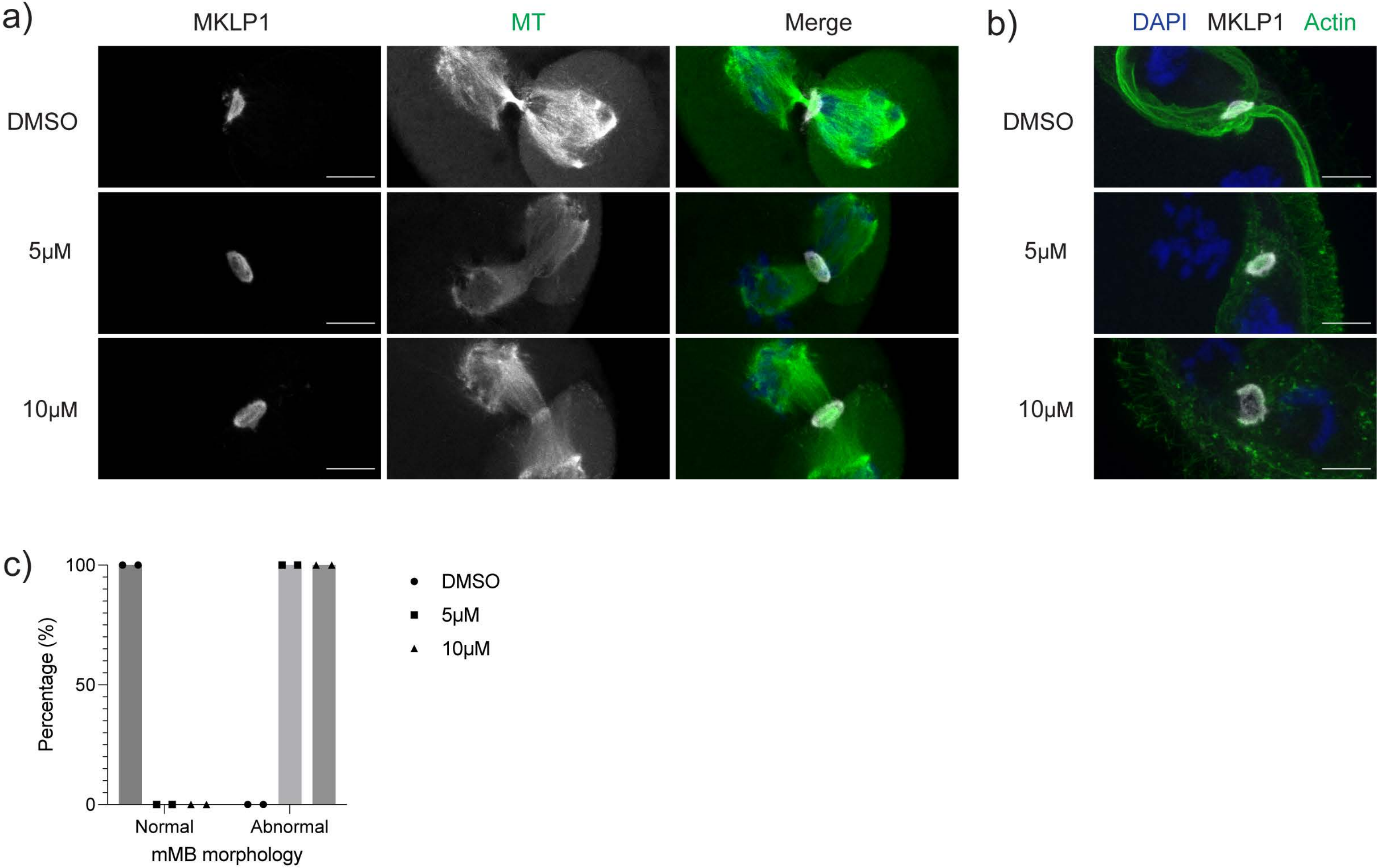

### Fig. S4

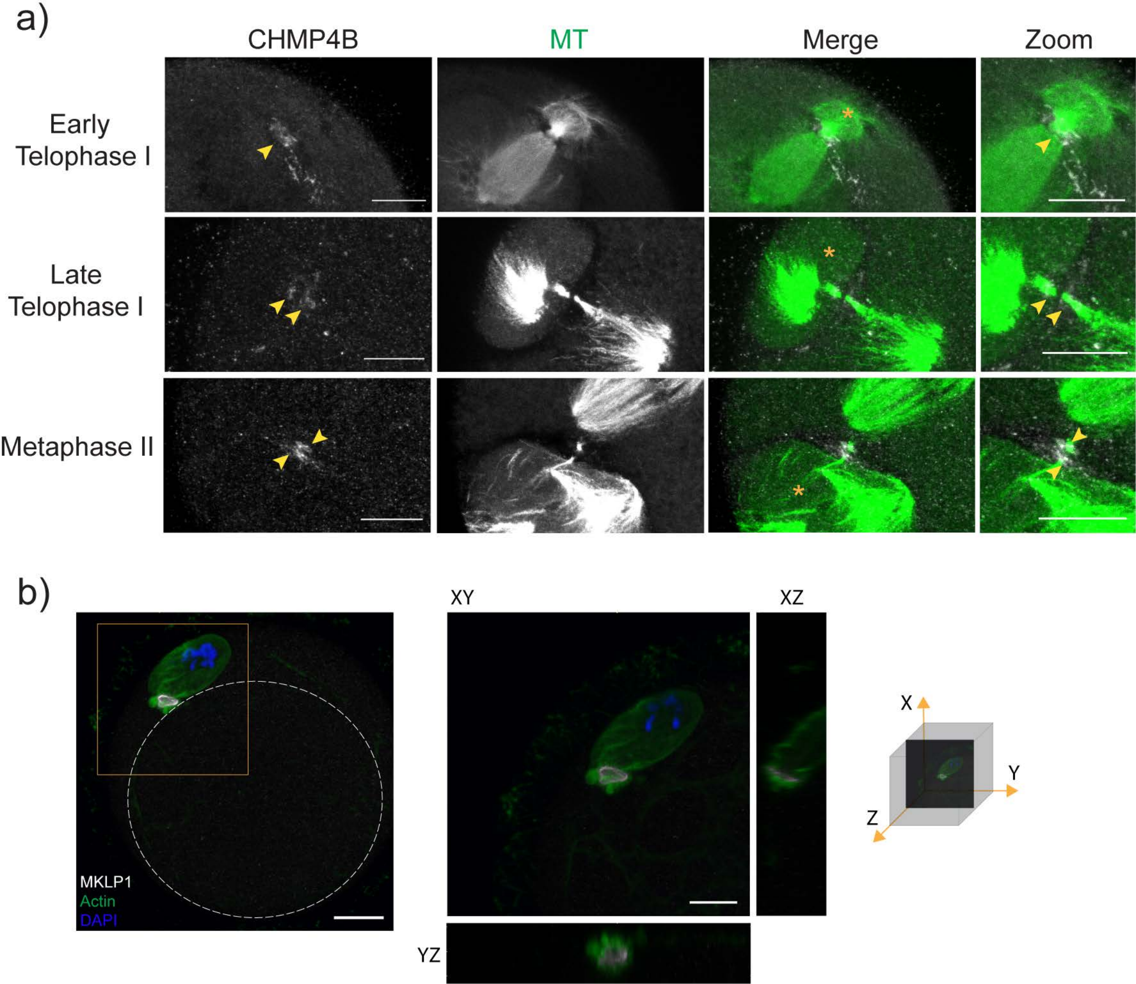

### Fig. S5

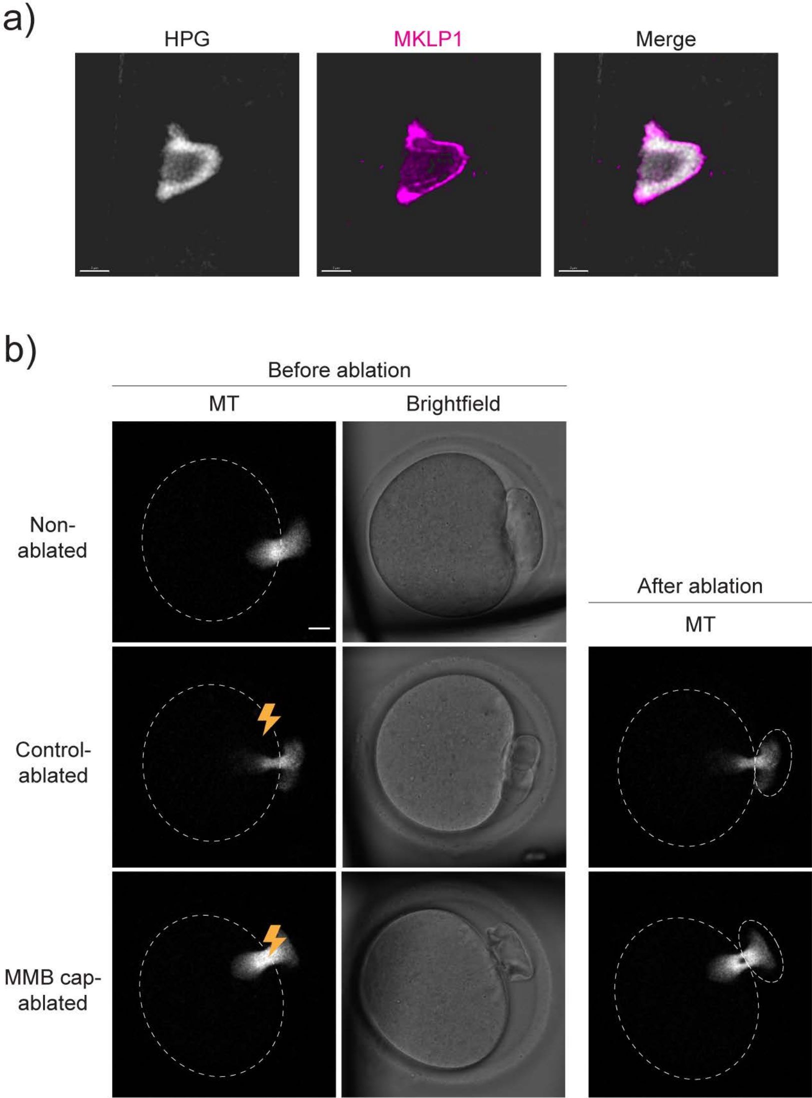
